# Housing Mice in Thermoneutrality Causes Tissue-specific Changes in Number, Identity, and Phase of Circadian-expressed mRNA Transcripts

**DOI:** 10.64898/2026.05.05.722706

**Authors:** Abhilash Prabhat, Shrishti Naidu, Isabel G. Stumpf, Emma J. Clemons, Samuel O. Nwadialo, Ezekiel Rozmus, Yuan Wen, Karyn A. Esser, Elizabeth A. Schroder, Brian P. Delisle

## Abstract

Most laboratory mice are housed at room temperature (20-25°C), which exposes them to chronic mild cold stress because it is below their thermoneutral temperature (30°C). We hypothesized that mild cold stress suppresses circadian gene expression in peripheral tissues. We performed RNA sequencing on hearts, livers, and diaphragms collected every 4 hours over 48 hours in constant darkness from male mice to identify transcripts with approximately 24-hour rhythms. Thermoneutral housing produced tissue-specific changes in the number, identity, and timing of rhythmic transcripts without altering the expression of core circadian clock genes. In the heart, the number of rhythmic transcripts increased fourfold, whereas the diaphragm showed a 1.5-fold increase. In the liver, the overall number of rhythmic transcripts showed little change, but their identity changed by 30%. Gene Ontology analysis revealed coordinated changes in the temporal organization of metabolic pathways in the heart and liver. Together, these findings demonstrate that ambient housing temperature is a major determinant of tissue-specific circadian gene expression, altering the abundance, identity, and timing of rhythmic transcripts independently of the core circadian clock.

**Significance:** Scientists typically house laboratory mice at room temperature, below their thermoneutrality, forcing them to increase their metabolic rate to maintain core body temperature. Since ambient temperature plays an important role in metabolism, cold stress could disrupt circadian gene expression. Comparing mice housed at room temperature with a warmer, thermoneutral temperature, we found that housing temperature causes tissue-specific differences in rhythmically expressed genes in the heart, liver, and diaphragm, without altering core clock genes. The heart was especially sensitive, with rhythmic genes peaking at the transition between subjective light and dark cycles, increasing fourfold. These results identify ambient housing temperature as an underrecognized variable that biases circadian gene expression in cardio-metabolic tissues, affecting interpretation of preclinical studies of metabolism and disease.

## Introduction

Laboratory mice are routinely housed at room temperature (20-25°C), which is below their thermoneutral temperature (TN, ≈30°C) (1). Under these mildly cold-stressed conditions, mice increase energy expenditure, food intake, and basal metabolic rate by 30-40% (2). Mild cold stress may alter hormonal and autonomic signaling, thereby affecting circadian gene expression in peripheral tissues (3–6). We tested the hypothesis that housing temperature substantially alters the circadian gene expression in the heart, liver, and diaphragm of male mice. We focused on the heart and diaphragm because their activity changes with metabolic demand, whereas the liver is a central regulator of whole-body metabolism (**Figure 1A**).

**Figure 1:**
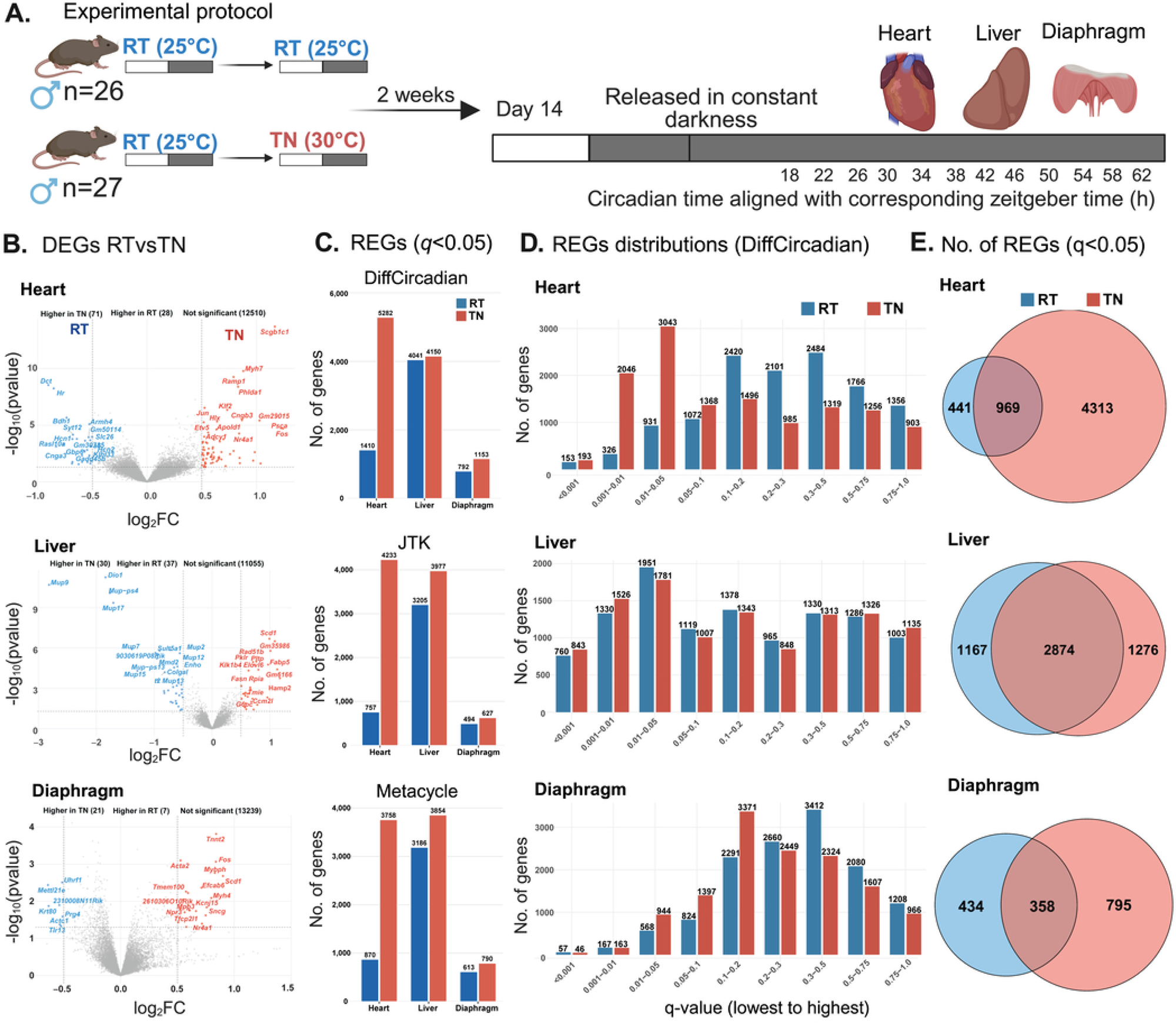
Thermoneutrality increases rhythmically expressed genes in the circadian cardio-metabolic transcriptome. **A.** Experimental protocol: Male SV129s/J mice (n = 53) housed at room temperature (RT; 25 ± 1 °C) in a 12-hour light/12-hour dark cycle with ad libitum access to food and water were assigned to one of two temperature conditions: continued RT (n = 26) or thermoneutrality (TN; 30 ± 1 °C, n = 27) for two weeks. After two weeks, mice were released into free-running, constant-dark conditions. Heart, liver, and diaphragm tissues were collected across the circadian cycle at 12 time points over 48 hours, starting at circadian time (CT) 18 aligned to the corresponding zeitgeber time; n = 2–3 mice/time point/housing condition. RNA sequencing was performed, and differentially expressed genes (DEGs) and rhythmically expressed genes (REGs) were identified in samples with more than 10 normalized counts/housing condition. **B.** Volcano plot showing upregulated DEGs under RT (blue) and TN (red) in heart, liver, and diaphragm tissues with non-significant genes (grey) at *p* < 0.05 and log2Fold change ± 0.5. **C.** Histograms showing the number of REGs using DiffCircadian, JTK-CYCLE, and MetaCycle statistical models under RT (blue bar) and TN (red bar) in heart, liver, and diaphragm tissues at *q* < 0.05. **D.** Histograms showing the number of REGs at different FDR-corrected *q* levels from <0.001 to 1.0 in mice under RT (blue bar) and TN (red bar) in heart, liver, and diaphragm tissues. **E.** Venn diagrams of the number of REGs in mice under RT (blue circle) and TN (red circle), and shared REGs in the heart, liver, and diaphragm tissues at *q* < 0.05.

## Results and Discussion

### Thermoneutral housing selectively alters rhythmic gene expression across tissues

Of 55,487 annotated genes, 12,609 in the heart, 11,122 in the liver, and 13,267 in the diaphragm met the expression thresholds (>10 normalized counts across all samples under both housing conditions) and were included in the analyses of differential expression and rhythmicity. Fewer than 1% of expressed transcripts differed in overall abundance between room temperature and thermoneutral housing in any tissue (**Figure 1B**).

To identify rhythmic transcripts, we applied three different methods, DiffCircadian, JTK-CYCLE, and MetaCycle (7–9), which produced similar results across tissues and across housing conditions at FDR-corrected *q* < 0.05 (**Figure 1C, D)**. The number of rhythmic transcripts detected in the hearts, livers, and diaphragms of mice housed at room temperature was consistent with previous studies of adult male mice (7–9).

Thermoneutral housing altered rhythmic gene expression in a tissue-specific manner (**Figure 1E**). In the heart, the proportion of rhythmic transcripts increased from 11% to 42%. In contrast, the liver showed little change in the overall proportion of rhythmic transcripts (36% vs. 37%), whereas the diaphragm showed a modest increase (6% to 9%). Despite the stable overall proportion of rhythmic liver transcripts, more than 30% of rhythmic transcripts differed between room temperature and thermoneutral housing, indicating substantial remodeling of the hepatic circadian transcriptome.

### Thermoneutral housing reshapes the timing of rhythmic gene expression

Compared with room temperature, thermoneutral housing produced a more concentrated distribution of transcript peak times in the heart and liver. In both tissues, rhythmic transcripts clustered near the end of the rest cycle (CT 8-12) and the end of the active cycle (CT 20-24), whereas the effect on the diaphragm showed a more complex pattern (**Figure 2A**).

**Figure 2:**
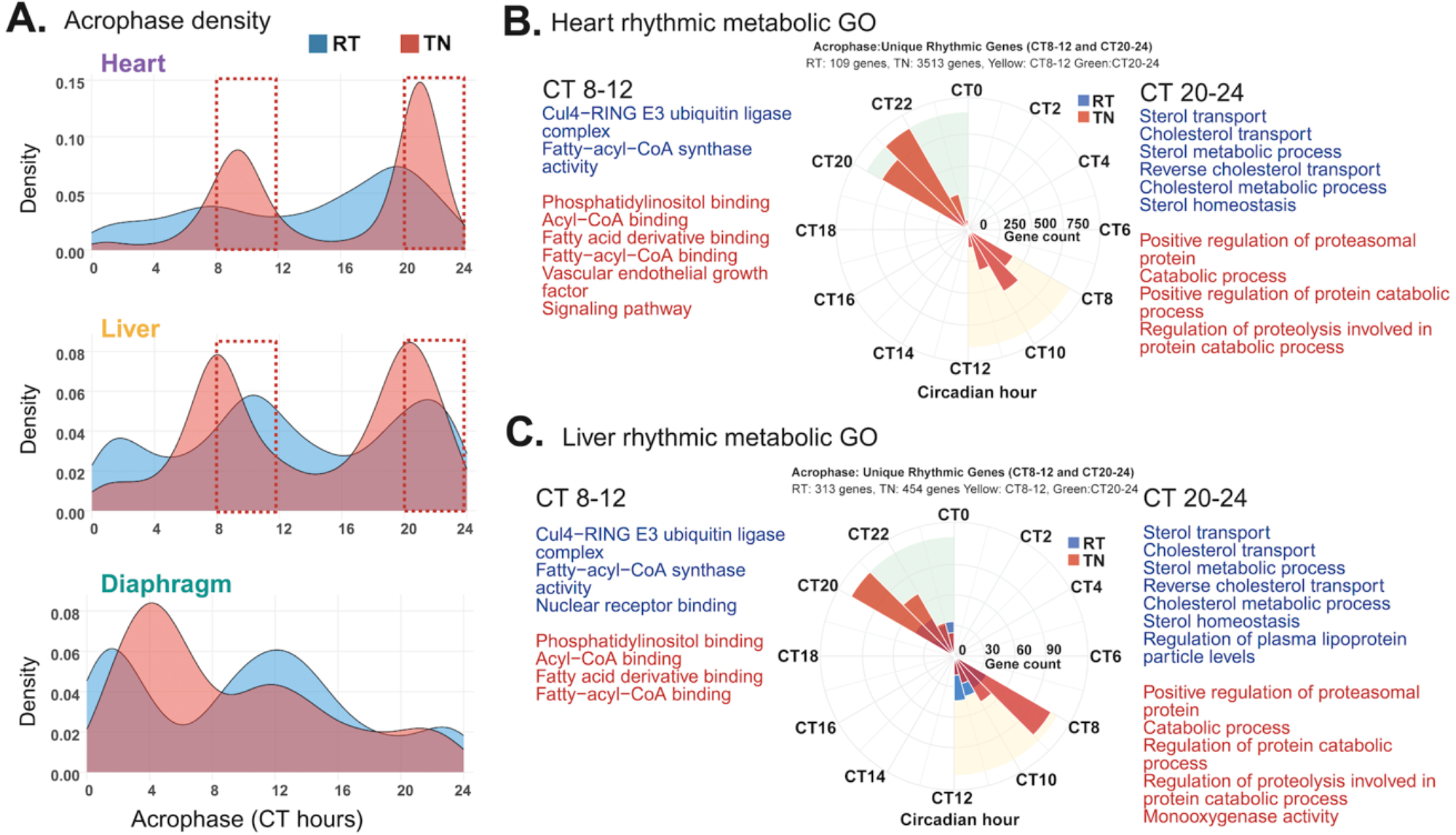
Thermoneutrality enriched time-dependent REGs and metabolic pathways. **A.** Acrophase density plot of rhythmically expressed genes (REGs) in mice under room temperature (RT; blue) and thermoneutrality (TN; red), in the heart, liver, and diaphragm at different circadian times (CT). The red dotted box represents higher acrophase density at CT 8-12 and CT 20-24 in heart and liver tissue. **B.** The circular plot shows unique REGs at CT 8-12 and CT 20-24 in the heart. The blue and red shading and font show RT- and TN-specific significant metabolic genome ontology (GO) terms (*q* < 0.05). **C.** The circular plot shows unique REGs at CT 8-12 and CT 20-24 in the liver. The blue and red shading and font show RT- and TN-specific significant metabolic genome ontology (GO) terms (*q* < 0.05).

In the heart, transcripts that peaked near the end of the rest cycle at room temperature were enriched for circadian rhythm and glucose-response pathways. In contrast, transcripts peaking at the same circadian phase under thermoneutral conditions were enriched for Ca2+ transport, ion channel regulation, and muscle contraction (**Dataset 1**). Rhythmic cardiac transcripts that peaked near the end of the active cycle under thermoneutral conditions were enriched for fatty acid oxidation and mitochondrial function, consistent with previous reports of circadian regulation of metabolic pathways in gastrocnemius skeletal muscle (10).

In the liver, transcripts that remained rhythmic under both housing conditions were enriched for similar biological pathways. Liver transcripts rhythmic only at room temperature and peaking near the end of the active cycle were enriched for cholesterol homeostasis and lipoprotein metabolism, whereas transcripts rhythmic only under thermoneutral conditions were enriched for protein degradation via the proteasome and mitochondrial function (**Dataset 1**).

Genes rhythmic only at room temperature and those rhythmic only under thermoneutral conditions also showed broadly similar functional enrichment in metabolic-specific pathways in both the heart and liver. Near the end of the rest cycle, room temperature-specific rhythmic genes were enriched for E3 ubiquitin ligase complex and fatty-acyl-CoA synthase activity. In contrast, thermoneutral-specific rhythmic genes were enriched for phosphatidylinositol binding, fatty acyl-CoA binding, acyl-CoA binding, and vascular endothelial growth factor signaling pathways. Near the end of the active cycle, room temperature-specific rhythmic genes were enriched for cholesterol homeostasis and cholesterol and sterol metabolic processes, whereas thermoneutral-specific rhythmic genes were enriched for regulation of protein ubiquitination and protein catabolic processes, etc (**Figure 2B, 2C; Dataset 1**).

### Thermoneutrality increases rhythmic gene expression shared across tissues

The number of rhythmically expressed genes shared across the heart, liver, and diaphragm nearly doubled in mice housed in thermoneutrality, increasing from 118 genes at room temperature to 232 genes (**Figure 3A**). Sixty-five rhythmic genes were conserved across all tissues and housing conditions and were enriched for rhythmic processes, circadian rhythm, regulation of circadian rhythms, circadian regulation of gene expression, regulation of myeloid cell differentiation, and intracellular receptor signaling pathways, etc (**Figure 3B**). These transcripts also showed similar phases across all three tissues (**Figure 3C**). Core circadian clock genes and clock-associated transcripts remained rhythmic under both housing conditions and showed similar mean expression levels (mesor), amplitudes, and phases (**Figure 3D; Dataset 1**).

**Figure 3:**
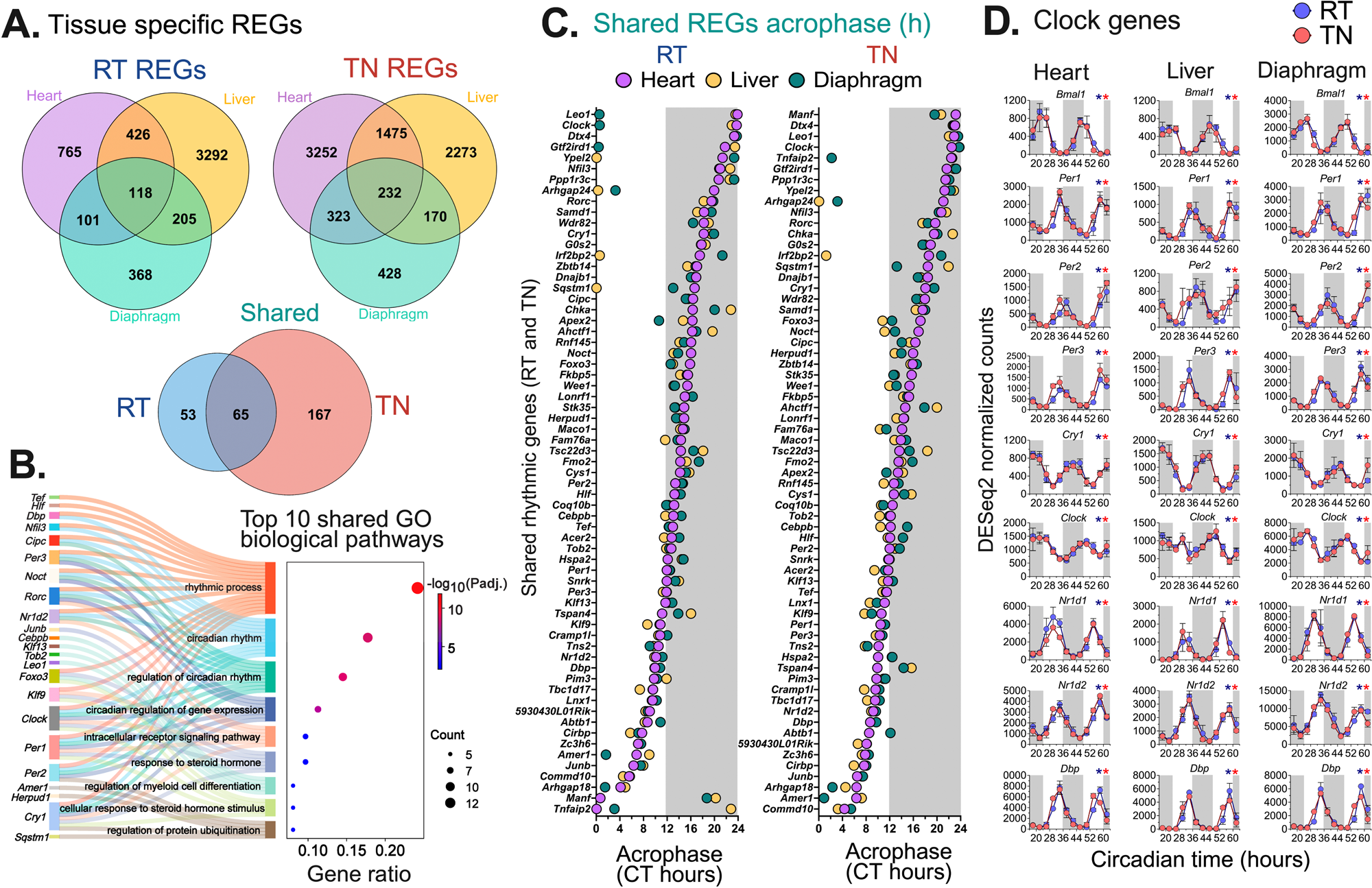
Thermoneutrality increases shared REGs across tissues and preserves clock gene expression pattern. **A.** Venn diagram of room temperature rhythmically expressed genes (RT-REGs), thermoneutrality (TN-REGs), and RT (blue circle) and TN (red circle) shared REGs across the heart (purple circle), liver (yellow circle), and diaphragm (green circle) tissues of mice at *q* < 0.05. **B.** The Sankey-alluvial plot shows the top 10 shared enriched GO biological pathways in all three tissues at *q* < 0.05, with gene names listed on the Y-axis. **C.** The acrophase of the heart (purple), liver (yellow), and diaphragm (green) tissue shared REGs under RT and TN. The Y-axis labels are the gene abbreviations. The unshaded and shaded regions represent subjective rest (CT0-12) and subjective active (CT12-24) hours. The acrophase of liver and diaphragm are aligned to the heart phase. **D.** Mean (SD) DESeq2 normalized counts of selected core clock genes plotted across circadian timepoints (CTs in hours) in heart, liver, and diaphragm tissues under RT (blue) and TN (red). The unshaded and shaded regions represent subjective rest (CT0- 12) and subjective active (CT12-24) hours. The blue and red asterisks indicate significant rhythmicity under RT or TN at *q* < 0.05.

The tissue-specific changes in rhythmic gene expression occurred without detectable changes in core circadian clock transcript expression, suggesting that ambient temperature modifies rhythmic transcription downstream of, or in parallel with, the core circadian clock. Consistent with this idea, decoupleR analysis (11) predicted greater cyclic AMP-responsive element modulator (Crem) activity at room temperature and greater Gata-binding protein 4 (Gata4) and ETS transcription factor ELK 4 (Elk4) activity at thermoneutrality in the heart (**Figure 4A**). Promoter motif analysis using HOMER identified enrichment of Klf4, ZEB2, Sp1, retinoic acid receptor gamma (RARg) and COUP-TFII motifs among genes rhythmic only in thermoneutrality, whereas genes rhythmic only at room temperature were enriched for Zfp281, RORg and Hoxa11 motifs. Genes rhythmic under both conditions were enriched for thyroid hormone receptor alpha/beta (THRa/b), PU.1-IRF, MafK and COUP-TFII motifs (**Figure 4B**). Integrating the HOMER and decoupleR analyses identified FOX-family and GATA:SCL binding motifs as candidate regulators of thermoneutrality-induced rhythmic gene expression (**Figure 4C**). Because *Foxa3* is not expressed in the adult mouse heart, the enriched forkhead motifs are most likely to reflect the activity of other FOX family transcription factors.

**Figure 4:**
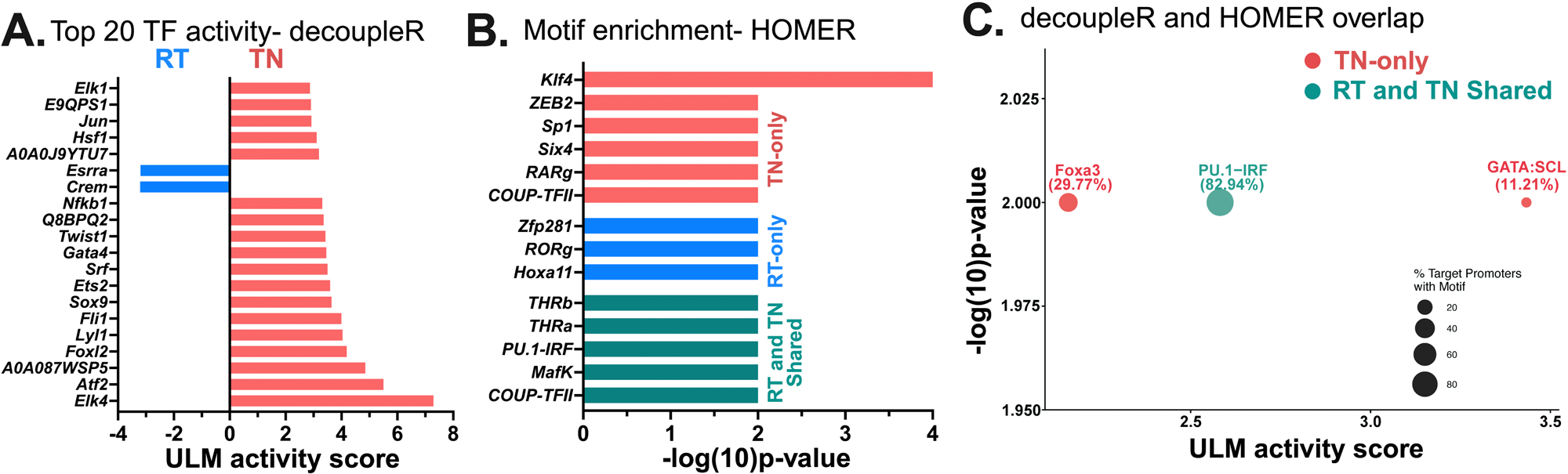
Thermoneutrality enhances distinct transcription factor activity in the heart. **B.** Histogram showing the top 20 significant transcription factor (TF) activity using the univariate linear model (ULM) in room temperature (RT; blue bar) and thermoneutrality (TN; red bar) by decoupleR in the cardiac transcriptome at *p* < 0.05. **C.** Histogram showing top significant motif-enriched TFs in RT-only (blue bar) and TN- only (red bar) and RT and TN-shared (green bar) rhythmically expressed genes (REGs) by HOMER at *p* < 0.05. **D.** Overlap of decoupleR and HOMER representing highly active TFs and motif- enriched TFs in cardiac REGs.

Our findings identify ambient housing temperature as a major regulator of rhythmic gene expression in peripheral tissues. Compared with thermoneutrality, which more closely reflects the thermal environment experienced by humans (12, 13), standard laboratory housing exposes mice to chronic mild cold stress that increases energy expenditure and metabolic rate by twofold (2). Under thermoneutral conditions, rhythmic gene expression was extensively reorganized in a tissue-specific manner, with the largest change occurring in the heart, where the number of rhythmic genes increases fourfold. In contrast, the number of rhythmic genes detected in different tissue types at room temperature was comparable to previous studies in male mice (7–9), indicating that thermoneutrality produces a distinct biological response rather than reflecting technical differences between studies.

Given the magnitude of these temperature-driven changes in rhythmic gene expression in cardio-metabolic tissues, it raised the question of whether temperature was acting on the clock mechanism itself or on its downstream outputs. Despite this extensive remodeling, thermoneutrality did not alter the expression level, amplitude, or phase of core clock genes in any tissues tested. This dissociation is consistent with previous work showing that the central pacemaker in the suprachiasmatic nucleus is resistant to temperature-resetting, whereas peripheral tissues respond to physiological temperature signals (14–16). The tissue-specific changes in the REGs were not secondary to changes in core body temperature because TN housing does not affect core body temperature in male mice (1, 2, 17), providing a possible explanation for the preserved circadian clock expression. Instead, ambient temperature appears to modify rhythmic gene expression through tissue-specific regulatory pathways. Consistent with this model, our integrated decoupleR and HOMER analyses (11) identified distinct transcriptional regulators, including Crem at room temperature and Gata4, Elk4, and FOX-family transcription factors at thermoneutrality.

The biological pathways associated with rhythmic genes also shifted with housing temperature in a time-dependent manner. In the heart, genes peaking near the transition from rest to activity were enriched for fatty acid oxidation and mitochondrial function at thermoneutrality, whereas room temperature preferentially enriched pathways involved in glucose handling. This pattern resembles the temporal partitioning of energy substrate use described in gastrocnemius skeletal muscle (10) and is consistent with circadian regulation of mitochondrial oxidative metabolism through rhythmic NAD+ biosynthesis (18). In the liver, thermoneutrality shifted rhythmic enrichment away from cholesterol and lipoprotein metabolism toward proteasomal degradation and mitochondrial pathways, consistent with previous reports that housing temperature reshapes hepatic circadian metabolism (4). Together, these findings suggest that ambient temperature modifies the timing of metabolic programs across multiple peripheral tissues.

The heart exhibited the largest increase in rhythmic gene expression despite an intact core circadian clock, highlighting the sensitivity of cardiac transcriptional programs to environmental temperature. The heart relies on a cell-autonomous circadian clock to anticipate predictable daily changes in workload and energy demand (19). Previous studies have also shown that thermoneutral housing alters autonomic regulation by increasing the vagal influence on the heart and slowing average heart rate by more than 17% in male mice (6). Thermoneutral housing also modifies autonomic signaling to the liver (5). These physiological adaptations provide a plausible mechanism by which ambient temperature could reshape peripheral rhythmic gene expression independent of or in parallel with the core circadian clock.

### Limitations

This study was limited to young adult male mice. Because sex and age both influence circadian gene expression (7, 20, 21), it will be important to determine whether these findings generalize across biological variables. Future studies should also define the time course required for thermoneutral remodeling, determine the range of temperatures that influence rhythmic gene expression, and establish whether similar responses occur in other tissues. Gene Ontology enrichment analyses are also influenced by the number of genes included in the analysis. Therefore, the broader range of enrichment pathways observed in the heart at thermoneutrality may partly reflect the larger number of rhythmic genes detected under this condition.

Overall, these findings identify housing temperature as an underappreciated determinant of peripheral circadian gene expression and demonstrate that standard laboratory housing conditions systematically reshape rhythmic transcription, particularly in the heart.

## Materials and methods

### Animals and housing conditions

All animal procedures complied with the Association for Assessment and Accreditation of Laboratory Animal Care guidelines and were approved by the Institutional Animal Care and Use Committee (Protocol number: 2019-3304) at the University of Kentucky. Male SV129s/J mice procured from the Jackson Laboratory (n=53) were housed at room temperature (25 ± 1 °C) in a 12-hour/light/12-hour dark cycle with ad libitum access to food and water. After one week of acclimation, mice were assigned to one of two temperature conditions: continued room temperature (25 °C ± 1, °C n=26) or thermoneutrality (30 ± 1 °C, n=27) for two weeks under ad libitum access to food and water. At the end of day 14, mice were released in free-running constant dark conditions (0 lx) to assess the endogenous circadian transcriptome independent of external light cues.

### Circadian sampling

Tissue collection was performed on day 15 following release into constant dark conditions. Animals were sampled across the circadian cycle at 12 time points over 48 hours (circadian time 18, 22, 26, 30, 34, 38, 42, 46, 50, 54, 58, and 62 hours aligned to the corresponding zeitgeber time), with n = 2–3 mice per time point.

### Tissue collection

Mice were quickly euthanized under dark conditions, and tissues were rapidly collected. Heart (ventricles), liver, and diaphragm were dissected, flash frozen in liquid nitrogen, and stored at −80 °C until further processing.

### RNA extraction, library preparation, RNA sequencing

Total RNA was isolated using Qiagen RNeasy Mini Fibrous Kit (ventricles and diaphragm) and Qiagen RNeasy Plus Mini Kit (liver) following the manufacturer’s instructions. Library preparation was done, and samples were sequenced on the Illumina® NovaseqX Plus platform (Illumina, California, USA). Read counts were normalized using DESeq2 for further analysis.

### Circadian rhythm analysis

Rhythmic gene expression was assessed using three different methods at FDR- corrected *q* < 0.05. DESeq2 counts were filtered for >10 counts per sample per condition to select highly expressed genes. We applied JTK-CYCLE, MetaCycle (8), and DiffCircadian (7) packages with cosinor analysis and LR_Rhythmicity implemented in RStudio. For final analysis, the DiffCircadian package was used (**Figure 1C-E**).

### Transcription factor activity (decoupleR) and promoter motif enrichment analysis in the heart (HOMER)

Transcription factor activity was inferred from the differential expression results using the decoupleR package (v1.4.0) in R(11). The Univariate Linear Model method was applied using the DoRothEA regulon for mouse (confidence levels A, B, and C) as the TF-gene interaction network. Homer was used for promoter motif enrichment analysis. To identify specific transcriptional regulators of thermoneutral-induced rhythmicity, results from decoupleR transcription factor activity inference and HOMER promoter motif enrichment were integrated.

### Statistical analysis

All statistical analyses were conducted using R (RStudio 4.4.3) and PRISM (GraphPad 11.0.2, Boston, Massachusetts, USA) unless specified otherwise. Rhythmically expressed genes were defined at FDR-corrected *q* < 0.05 (Benjamini-Hochberg correction) values. Detailed statistical methods and thresholds are indicated in the corresponding methods sections and figure legends. All plots were generated in R using ggplot2, patchwork, and ggseqlogo packages and PRISM. Transcription factor binding motif sequence logos were generated from JASPAR2020 position weight matrices using ggseqlogo. No new code was generated for this manuscript.

## Data availability

Extended analyses and datasets are deposited in Figshare (DOI:10.6084/m9.figshare.32172870) and submitted with GEO accession number GSE337330. Details are in the Supplementary Information Appendix (*SI Appendix*).

## Acknowledgements

National Heart, Lung, and Blood Institute R01HL172813 to B.P.D. and E.A.S., Pathway to Independence Grant from Diabetes and Obesity Research Priority Area, and the Barnstable Brown Diabetes and Obesity Center, University of Kentucky, to A.P., National Institute of Arthritis and Musculoskeletal and Skin Diseases R00AR081367 to Y.W., and National Institute of Arthritis and Musculoskeletal and Skin Diseases R01AR079220 and National Institute on Aging P30AG028740 to K.A.E. The content is solely the responsibility of the authors and does not necessarily represent the official views of the NIH. Claude AI (Sonnet 4.6) was used for formatting and copyediting purposes.

