## Supplementary Information for "Housing Mice in Thermoneutrality Causes Tissue-specific Changes in Number, Identity, and Phase of Circadian-expressed mRNA Transcripts"

**Materials and methods**

*Animals and housing conditions*

All animal procedures complied with the Association for Assessment and Accreditation of Laboratory Animal Care guidelines and were approved by the Institutional Animal Care and Use Committee (Protocol number: 2019-3304) at the University of Kentucky. Male SV129s/J mice procured from the Jackson Laboratory (n = 53) were housed at room temperature (25 ± 1 °C) in a 12-hour/light/12-hour dark cycle (LD) with ad libitum access to food and water. After one week of acclimation, mice were assigned to one of two temperature conditions: continued RT (25 °C, n=26) or thermoneutrality (TN; 30 ± 1 °C, n=27) for two weeks under ad libitum access to food and water. At the end of day 14, mice were released in free-running constant dark conditions (DD; 0 lx) to assess the endogenous circadian transcriptome independent of external light cues.

*Circadian sampling*

Tissue collection was performed on day 15 following release into DD. Animals were sampled across the circadian cycle at 12 time points over 48 hours (circadian time 18, 22, 26, 30, 34, 38, 42, 46, 50, 54, 58, and 62 hours aligned to the corresponding zeitgeber time), with n = 2–3 mice per time point. Circadian time was determined based on the prior LD schedule. As in our previously validated circadian transcriptomic studies (1, 2), n = 2–3 mice per time point were employed, with rhythmicity detection power derived from the 12-timepoint, 48-hour temporal structure and the cosinor-based framework of DiffCircadian.

*Tissue collection*

Mice were quickly euthanized under dark conditions, and tissues were rapidly collected. Heart (ventricles), liver, and diaphragm were dissected, flash frozen in liquid nitrogen, and stored at −80 °C until further processing.

*RNA extraction, library preparation, RNA sequencing*

Total RNA was isolated using Qiagen RNeasy Mini Fibrous Kit (ventricles and diaphragm) and Qiagen RNeasy Plus Mini Kit (liver) following the manufacturer’s instructions. Isolated RNA sample quality was assessed by High Sensitivity RNA Tapestation (Agilent Technologies Inc., California, USA) and quantified by AccuBlue® Broad Range RNA Quantitation assay (Biotium, California, USA). Paramagnetic beads coupled with oligo d(T)25 are combined with total RNA to isolate poly(A)+ transcripts based on the NEBNext® Poly(A) mRNA Magnetic Isolation Module manual (New England BioLabs Inc., Massachusetts, USA). Prior to first-strand synthesis, samples are randomly primed and fragmented based on the manufacturer’s recommendations. The first strand is synthesized with the Protoscript II Reverse Transcriptase with a longer extension period, approximately 40 minutes at 42⁰C. All remaining steps for library construction were used according to the NEBNext® Ultra™ II Directional RNA Library Prep Kit for Illumina® (New England BioLabs Inc., Massachusetts, USA). The final library quantity was assessed by Qubit 2.0 (ThermoFisher, Massachusetts, USA), and quality was assessed by TapeStation D1000 ScreenTape (Agilent Technologies Inc., California, USA). Diaphragm samples yielded a low RIN score; hence, total RNA sequencing was applied. Final library size was about 430bp with an insert size of about 300bp. Illumina® 8-nt dual-indices were used. Equimolar pooling of libraries was performed based on QC values and sequenced on an Illumina® NovaseqX plus platform (Illumina, California, USA) with a read length configuration of 150 PE for 40M PE reads per sample (20M in each direction).

FastQC (version v0.12.1) was employed to check the quality of raw reads. Trimmomatic (version v0.39) was applied to cut adaptors and trim low-quality bases with the default settings. STAR package (version 2.7.10b) was used to align the reads via the mouse GRCm38 (*Mus musculus*, mm10) genome reference. The Picard tools (version 3.0.0) were used to mark duplicates in the mapping. StringTie (version 2.2.1) was used to assemble the RNA-Seq alignments into potential transcripts. HTSeq (version 2.0.3) was used to count mapped reads for genomic features such as genes, exons, promoters, gene bodies, genomic bins, and chromosomal locations. DESeq2 provided the normalized counts for further analysis.

*Circadian analysis*

Rhythmic gene expression was assessed using three different methods. We applied JTK-CYCLE, MetaCycle(3), and DiffCircadian(2) packages with cosinor analysis and LR_Rhythmicity implemented in RStudio. For final analysis, the DiffCircadian package was used (**Figure 1C-E**). Circadian rhythmicity was determined across the time series for each tissue and temperature condition. Genes were considered rhythmically expressed if they met a false discovery rate (FDR; Benjamini-Hochberg correction) threshold of *q* < 0.05 in each condition. ClusterProfiler (RStudio) gave genome ontology (GO) pathways. Time-point-specific REGs at the end of the rest (CT 8-12) and end of the activity (CT 20-24) periods were filtered for heart- and liver-specific REGs. Circadian, tissue-specific, and metabolic enriched GO were generated for the end of the rest (CT 8-12) and end of the activity (CT 20-24) periods, REGs at *pajdusted* < 0.05.

*Transcription Factor and transcription factor activity (decoupleR)*

Transcription factor (TF) activity was inferred from the differential expression results using the decoupleR package (v1.4.0) in R(4). The Univariate Linear Model (ULM) method was applied using the DoRothEA regulon for mouse (confidence levels A, B, and C) as the TF-gene interaction network. Activity scores were computed for all expressed genes in the heart, with positive scores indicating higher TF activity in TN and negative scores indicating higher activity in RT. TFs with a *p* < 0.05 were considered statistically significant. To characterize TF regulatory programs within each rhythmicity group, significant TF target genes were intersected with each gain of rhythm genes (GORG) group (TN-only, RT-only, rhythmic in both) and ranked by number of targets and consistency of regulation direction with TF activity score (**Dataset 1**). For all the analyses, *p* < 0.05 and *p-adjusted* < 0.05 are reported.

*Promoter motif enrichment analysis in the heart (HOMER)*

We classified genes into four groups based on rhythmicity: genes gaining rhythmicity in TN (TN-only rhythmic), genes losing rhythmicity in TN (RT-only rhythmic), genes rhythmic under both conditions, and non-rhythmic genes under either condition using FDR *q* < 0.05 (**Figure 1D: Heart**). Differential expression between TN and RT conditions was assessed using limma-voom, and the results were integrated with rhythmicity classifications to characterize the transcriptional landscape of each gene group. Promoter regions (−2,000 to +200 bp relative to the transcription start site) were extracted for each GORG group using the TxDb.Mmusculus.UCSC.mm10.knownGene and BSgenome.Mmusculus.UCSC.mm10 Bioconductor packages in R. Gene symbols were mapped to Entrez IDs using the org.Mm.eg.db annotation package. Genomic coordinates were exported as BED files for use with HOMER (Hypergeometric Optimization of Motif EnRichment, v5.1). Known motif enrichment analysis was performed using findMotifsGenome.pl with the mm10 genome, using all expressed genes (n=11,864 promoter regions) as background. Motif lengths of 8, 10, and 12 bp were tested using 4 parallel processors (-p 4 -len 8,10,12 -size given). Statistical significance was assessed by HOMER's built-in binomial/hypergeometric test with Benjamini-Hochberg correction for multiple comparisons. Motifs with *p* < 0.05 were considered significant; motifs with *q* < 0.05 were considered significant after multiple testing correction.

*Overlapping transcription factor activity with HOMER*

To identify specific transcriptional regulators of thermoneutral-induced rhythmicity, results from decoupleR TF activity inference and HOMER promoter motif enrichment were integrated. TF details from HOMER output were mapped to their corresponding gene symbols using a curated synonym dictionary accounting for naming conventions across databases (e.g., GATA:SCL with Gata4, PU.1-IRF with Spi1, and Foxa3 with Foxa2 family). FOXA2 and FOXA3 share ~95% sequence identity in their DNA-binding domain (forkhead domain). TFs identified as significant in both analyses; elevated activity score in decoupleR (*p* < 0.05, ULM) and enriched binding motif in GORG promoters (*p* < 0.05, HOMER) were plotted and classified as candidate regulators.

*Statistical analysis*

All statistical analyses were conducted using R (RStudio 4.4.3) and PRISM (GraphPad 11.0.2, Boston, Massachusetts, USA) unless specified otherwise. DESeq2 normalized expression values were used for differential expression analysis. DiffCircadian (RStudio) was used to identify rhythmically expressed genes at FDR-corrected *q*<0.05 (Benjamini-Hochberg correction) values with cosinor to accurately assess the mesor, amplitude, and acrophase (time of the amplitude). Detailed statistical methods and thresholds are indicated in the corresponding methods sections and figure legends. All plots were generated in R (v4.4.3) using ggplot2, patchwork, and ggseqlogo packages and PRISM. TF binding motif sequence logos were generated from JASPAR2020 position weight matrices using ggseqlogo. No new code was generated for this manuscript.
